# Automated and rapid self-report of nociception in transgenic mice

**DOI:** 10.1101/732305

**Authors:** Christopher J. Black, Anusha B. Allawala, Kiernan Bloye, Kevin N. Vanent, Muhammad M. Edhi, Carl Y. Saab, David A. Borton

## Abstract

A time-resolved, conscious report of detected nociceptive stimuli in mice offers an opportunity to examine the relationship between higher order neural circuits and pain perception. We have developed a detection behavior in transgenic mice that leverages temporally precise and cell-specific stimulation to elicit self-reports of nociception. Conscious reporting of peripheral nociceptive input may help identify neural mechanisms that generate pain perception.

## Main

Rodents are the predominate animal model for elucidating the neural mechanisms underlying pain; reflexive and avoidance behaviors are commonly used to evaluate peripheral and spinal pain processing^1^. Both genetic manipulations^2^ and *in vivo* neural recording methods^3^ have been widely used to examine the neural contributions to hypersensitivity. However, due to the temporal imprecision in delivering nociceptive stimuli, and the inherent ambiguity of interpreting behavioral reports, prior efforts have had limited success in mapping the neural circuits underlying nociceptive processing with fine temporal precision and high cellular specificity.

Nociceptive reflex behaviors have been shown to positively correlate with self-reports of pain in humans^4^. However, noxious stimulation can elicit spinal reflexes without recruiting higher order neural circuits in the brain^5^. Additionally, the classification of reflex behaviors is observer-dependent, thus potentially biasing the interpretation of the collected neurophysiological data. Alternatively, cognitive processes can be inferred from operant behaviors, for example learned aversion^1^. Such high-order processes, however, fail to capture the millisecond dynamics of decision-making and pain perception.

Accounting for cognitive processes is complicated by the fact that the instruments used to evoke pain are not nociceptive-specific^6^. Probing the skin via Von Frey filaments or radiant heat may also activate innocuous mechano- or thermo-receptors^7^. Posture itself can further influence the outcome of nociceptive stimulation; the positioning of limbs and digits has been shown to modulate nociceptive reflexes in rodents^6^ and the activity of touch-evoked cortical neurons in non-human primates^8^. Ultimately, nociceptive stimuli lack the temporal precision necessary to properly synchronize the millisecond scale neural activity to the evoked behavioral observation.

Therefore, we have developed an observer-independent, fast and temporally-precise lick-based detection task using peripheral optogenetic stimulation of A-delta and C-fiber afferents in transgenic mice.

To achieve A-delta and C-fiber specific stimulation, we used transgenic mice that express the light-sensitive ion-channel channel-rhodopsin2 (ChR2) in transient receptor potential V1 (TRPV1) containing neurons, which are responsible for peripheral transmission of noxious heat^9^. It has been shown previously that optogenetic activation of the hindpaw glabrous skin in this TRPV1-ChR2 mouse line specifically evokes electrophysiological responses in A-delta and C-fibers, and nocifensive behaviors without overt tissue damage^6,10,11^. We first performed immunohistochemical analysis to confirm ChR2 localization to small diameter nociceptors in the spinal cord and plantar surface of the hind paw tissue in these transgenic mice (**Fig.1a**). We then demonstrated that 470nm optogenetic stimulation of TRPV1-ChR2 mouse glabrous skin activated small diameter nociceptive sensory afferent neurons by performing acute, anesthetized, spinal extracellular electrophysiology in TRPV1-ChR2 mice (**Fig. 1b**). Conduction velocities computed from early (1.99±1.01 m/s) and late (0.20±0.01 m/s) evoked responses were in agreement with previously published conduction velocities in mouse A-delta and C-fiber primary nociceptors^12^.

**Figure 1.**
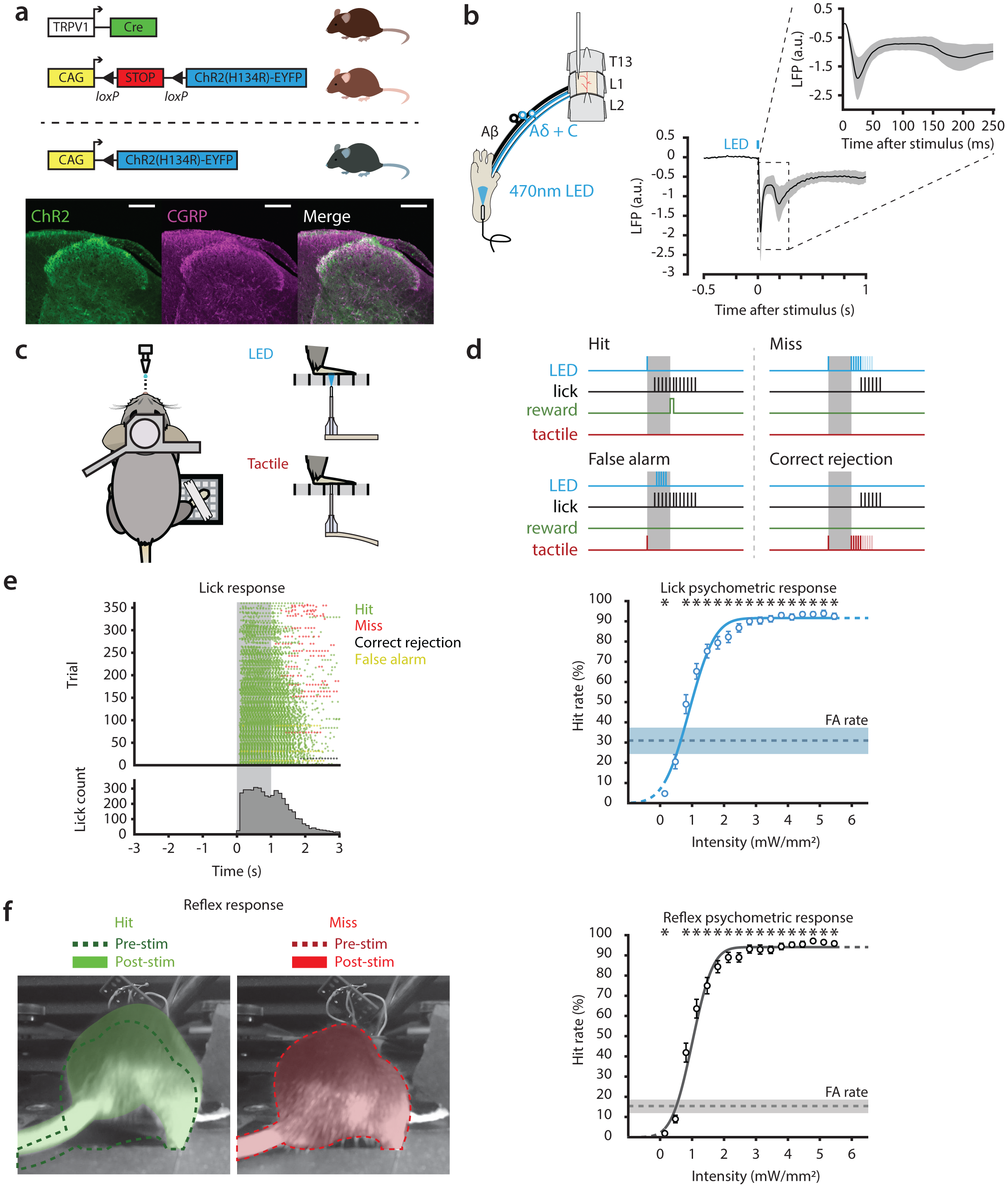
Validation of TRPV1-ChR2 and Dection Behavior. **(a)** (Top) Using a Cre driver line, we bred transgenic mice that expressed ChR2 in TRPV1 containing neurons. (Bottom) Immunohistochemistry of spinal cord slices of these TRPV1-ChR2 shows overlap of TRPV1 (green) with the nociceptive marker CGRP (magenta); scale bar = 150µm. **(b)** Acute, anesthetized spinal electrophysiology preparation (left) showing vertebral levels, where compound action potentials (CAP) were recorded during 10ms, 470nm LED pules to the hindpaw of TRPV1-ChR2 mice. The CAP waveform average (right, n = 6 mice, mean ± S.E.M.) shows two prominant post-stimulus peaks (t = 0) indicative of Adelta and C-fiber afferent recruitment; tick marks indicate 200ms. **(c)** Schematic layout of behavioral platform. Mice are headposted with direct access to a lick spout, and their right hindpaw is restrained over a grated floor. Optogenetic (470nm LED) and tactile stimuli are delivered through the grated floor at the same position underneath the right hindpaw. **(d)** Trial and response types; mice could either receive optogenetic (target) stimulation with a hit occuring when the mouse licks within the 1s report window and a miss occuring when the mouse does not lick within the 1s report window, or tactile (catch) stimulation with a false alarm occuring when the mouse licks within the 1s report window and a correct rejection occuring when the mouse licks during the 1s report window. **(e)** Lick raster and histogram (left, n = 1 session), grey shading indicates the 1s response window. Animals learned to lick to the onset of the optogenetic stimulus (green points) during the response window, with only a small percentage of trials resulting in not licking within the response window (red points). Cumulative psychometric responses for lick report (right) fitted with a cumulative Gaussian function (psignifit matlab toolbox) compared to false alarm (FA) rate (dotted line). **(f)** Reflexive report (left) showing two different trials that elicit a response (left, shaded green) and no response post-stimulus (right, shaded red) as compared to baseline position (dotted lines). Cumulative psychometric responses for reflex report (right) fitted with a cumulative Gaussian function (psignifit matla toolbox), compared to FA rate (dotted line). All psychometric responses showing mean ± SEM, n = 40 sessions from 5 mice. Asterisks in **e** and **f** denote significance compared to the respective false alarm rate, *P* < 0.025 using a two-sided Wilcoxon signed rank test, tested for multiple comparisons with Bonferroni correction.

We then trained the TRPV1-ChR2 mice to report detection of hindpaw optogenetic (470nm) stimulation by licking a water spout. TRPV1-ChR2 mice were head-fixed on a custom behavioral platform and were water restricted to incentivize lick reporting^13^. Mice were hindpaw restrained (**Fig. S1**) to ensure reproducibility of stimulus delivery across trials and sessions^14^. A ceramic ferrule attached to a fiber optic cable and piezoelectric bender provided optogenetic (target) and tactile (catch) stimulation to the hindpaw through a grated floor (**Fig. 1c**).

Mice received target stimuli consisting of a 10ms pulse of 470nm optogenetic stimulation at 17 intensities between 0.14-5.45 mW/mm2, and catch stimuli, consisting of a single tap by the ferrule (single cycle of a ∼120Hz sine wave activating mechanoreceptors). There were 20 trials for all 18 stimuli types (360 trials per session) that were delivered in a random order at 3-6s spaced inter stimulus intervals (ISI). During both target and catch trials, mice received an initial stimulus followed by a 1s response period (**Fig. 1d**). During target trials, licking within the response window was considered a hit, and mice had an 80% chance of receiving a calibrated water reward (5-10uL). Not licking during the response window was considered a miss, and mice received up to 10 additional stimuli (10ms pulse width, 100ms ISI). If mice licked following a catch stimulus (false alarm), they immediately received 5 ‘warning’ pulses of optogenetic stimulation (10ms pulse width, 100ms ISI, 5.45 mW/mm^2^). Mice that licked prior to the onset of the trial were given a time-out (3-6s). Sessions were only considered if the animal was engaged in the behavioral task, which was determined by a high sensitivity (d’ > 1.0) measured across the first 250 trials as well as the overall session.

Mice learned to report detection of target stimulation to the hindpaw with no significant responses elicited during 595nm control stimulation between 0.14-5.45 mW/mm2 (**Fig. S2**) indicating that lick responses were initiated by direct activation of ChR2 containing neurons in the glabrous skin. The animal’s hit rates were dependent on stimulus intensity, with responses reaching an asymptotic hit rate over 90% (**Fig. 1e**). Furthermore, the false alarm rate was significantly lower than the hit rates for stimuli around threshold level intensity. This indicates that mice learned to distinguish between nociceptive and tactile stimuli.

We then sought to compare the lick report to a standard withdrawal response. We recorded video data during detection behavior and scored reflexive responses elicited by optogenetic stimulation (**Fig. 1f**) that occurred within 250ms of stimulus onset. Scorers manually assigned values of 0 (no movement), 1 (mild movement), 2 (moderate movement), and 3 (overt movement) for each individual trial. Scores were then categorized as either a hit or miss by grouping “2” and “3” scores, and “0” and “1” scores, respectively. Similar to the lick report, the hit rate of the reflex response was dependent on stimulus intensity and reached hit rates over 90% in response to salient nociceptive stimuli. When we compared the reports at perceptual threshold, we found that both extrapolated threshold intensities (**Fig. S3**) and hit rates near perceptual threshold between lick and reflex reports were not significantly different (**Fig. 2a**, right). However, the lick report elicited a larger false alarm rate (30.9±21.0%) than the reflex response (15.2±17.2%).

**Figure 2.**
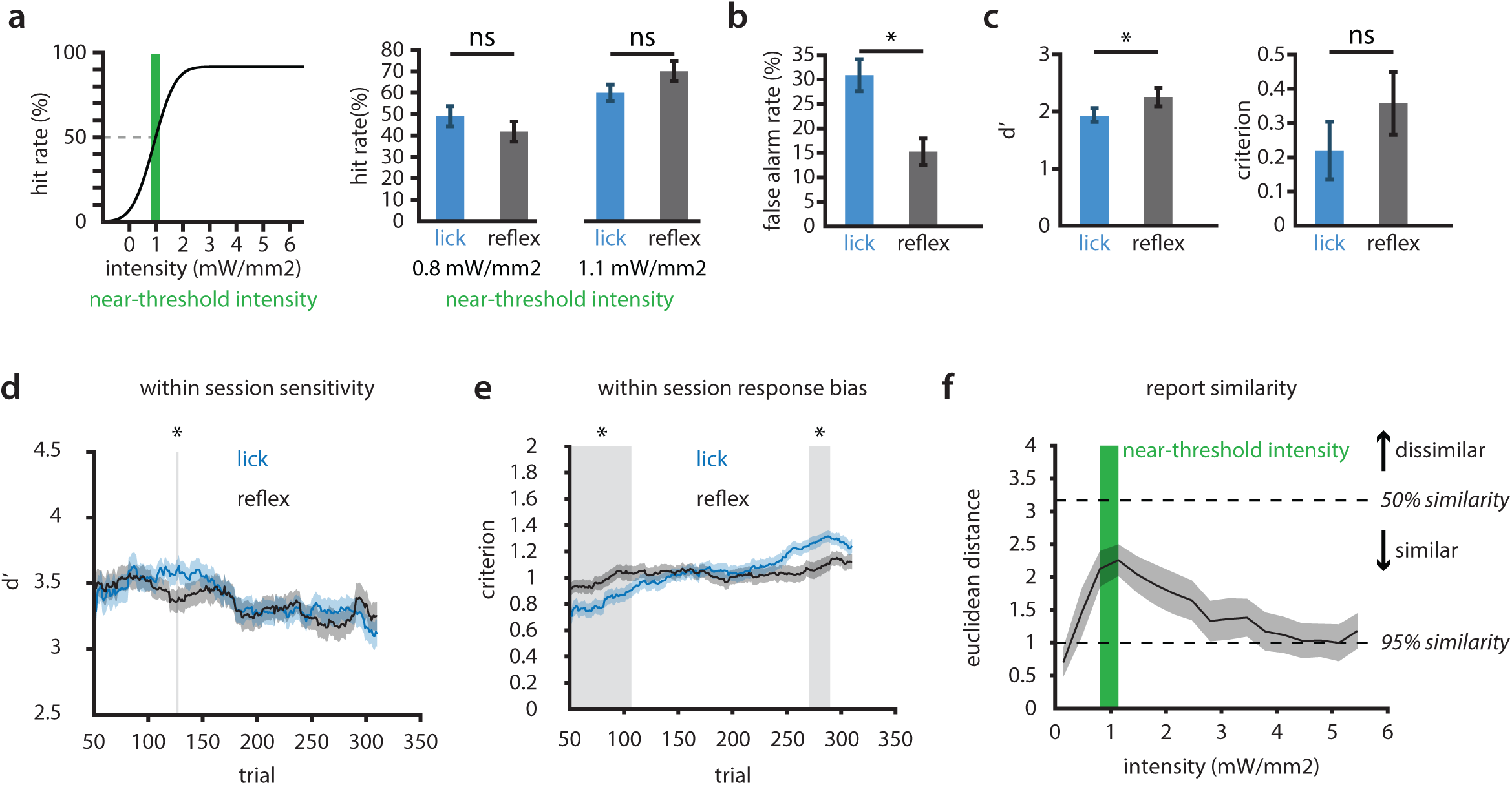
Comparison of lick-report and reflex withdrawal. **(a)** (Left) Illustration of near-threshold intensities taken from psychometric curve, (right) comparison of hit rate of lick (blue) and reflex (grey) reports at near-threshold intensities. **(b)** Comparison of false alarm rate, **(c)** sensitivity (d’), and response bias (criterion) between lick (blue) and reflex (grey) reports. **(d)** Comparison of sensitivity, and **(e)** response bias as a function of time for both lick and reflex reports. **(f)** Euclidean distance calculated as a function of stimulus intensity between lick and reflex. Green region indicates near-threshold intensities, while black dotted lines represent the euclidean distance for if response types were similar for 50% and 95% of all trials at any given intensity. All results are shown as mean±SEM, except for Euclide- an distance, which is shown as mean±95% CI. n = 40, grey shaded regions and asterisks in **a**-**e** indicate significance at *P* < 0.025 using a two-sided Wilcoxon signed rank test, **b** was tested for multiple comparisons with Bonferroni correction.

To determine differences in detection performance between the lick and reflex reports across all sessions, we calculated the sensitivity (d’) and response bias (criterion). The sensitivity of the reflex report was significantly greater across all sessions than the sensitivity for the lick report (**Fig. 2c**). The difference in sensitivity was expected, as the false alarm rate for the reflex report was significantly less than the false alarm rate for the lick report. The response bias, however, was not significantly different between the two reports (**Fig. 2c**).

We then looked at whether the sensitivity and response bias differed between the reports across time to determine if session averages masked any differences in detection performance. Therefore, we calculated the d’ and criterion in a sliding 50-trial window within sessions. The sensitivity of the lick and reflex withdrawal followed the same trajectory within sessions, which generated little difference between the reports (**Fig. 2d**). However, the lick criterion steadily increased as the reflex criterion remained constant (**Fig. S6**), which made the criterion of the reports significantly different in the beginning and the end of the sessions (**Fig. 2e**). Although additional investigation is necessary to determine the exact cause of the divergence in criterion; modulation of response bias has been attributed to the influence of top-down processes, such as motivation^15^, attention, and expectation, on perception and corresponding neural activity^16–18^. Finally, we wanted to determine if the reports differed on a trial-by-trial basis. We hypothesized that if the reflex report provided the same information as the lick report, then the intensity of the stimulus should not influence the similarity between the two reports. To test this, we calculated the Euclidean distance for each intensity across all sessions (**Fig. 2f**). We found that responses near perceptual threshold were significantly less similar (*P* < 0.025, two-sided Wilcoxon signed rank) than both sub-threshold and highly-salient, supra-threshold responses.

We have developed a method that achieves temporally precise, nociceptive-specific, self-reports over a large number of trials of a nocifensive stimulus in TRPV1-ChR2 transgenic mice. Our detection behavior is ideal for implementation with chronic neural recording techniques to study the circuit level mechanisms and dynamics that give rise to pain perception. The increasing response bias of the lick report within sessions might support the claims of strong top-down modulation^15,16,18^ in the mouse’s decision to report pain. Furthermore, the reduced similarity of the lick and reflex reports near-threshold intensities indicates that the lick and reflex reports might capture different features of the pain experience. Although TRPV1 expressing primary nociceptors make up a fraction of the total primary nociceptors, and therefore elicit only one modality of pain percept, we believe this platform can be expanded to other transgenic models, such as the SNS-ChR2 line that expresses ChR2 in alpha-SNS containing nociceptive dorsal root ganglion neurons^19^, to characterize the behavioral phenotypes and the underlying neural circuits of various pain states.

## Methods

### Animals

All animal experiments were approved by the Institutional Animal Care and Use Committee at Rhode Island Hospital (RIH), Providence RI. All mice, aged 2-24 months, were TRPV1-ChR2, bred at RIH using the TRPV1-Cre and Ai32(RCL-ChR2(H134R)/EYFP) commercially available lines from Jackson Laboratories. Mice (n = 6) in acute electrophysiology experiments were half female and half male, while mice used for behavior (n = 5) were male. Mice were housed in groups of two or more, unless they received a headpost implant, in which case they were housed in single cages.

### Headpost surgeries

Mice were anesthetized using Isoflurane. Their heads were shaved and sterilized using betadine and 70% ethanol. Mice were then fixed in a stereotactic frame (Kopf instruments). An incision was made down the midline of the scalp and the skin was retracted. The exposed skull was then cleaned and scored to provide a dry surface for bonding with the dental cement. Titanium or stainless-steel head posts were then properly aligned and fixed to the skull using dental cement (MetaBond). Mice were then placed under a heating lamp or on a warming pad until they were bright, alert, and responsive. Behavioral testing and water restriction began no less than one week following implantation.

### Water restriction

Mice were acclimated over a five-day period to water restriction. Mice were kept at or above 85% of their pre-restriction baseline weight. Daily water volume was determined based on this criterion but could be no less than 1 mL/day. Weight and body conditioning score were taken a minimum of 3 days per week and on every day mice performed behavior to assess their hydration status.

### Acute spinal electrophysiology

TRPV1-ChR2 mice were anesthetized with isoflurane and shaved from their hips to below the base of their neck. A 2cm incision was made on the surface of the back along the top of the spine. Following retraction of muscles along the spine, a laminectomy was performed on one vertebra between vertebral level T13 and L3. The vertebrae rostral to the laminectomy was clamped and raised to provide stability during recordings. Once the spine was stable, a durotomy was performed at the site of the laminectomy. A single epoxy coated, tungsten electrode (impedance 5MOhm, 127µm diameter, A-M Systems) was lowered into the dorsal horn of the spinal cord using a micromanipulator arm. General receptive fields were found first by tactile stimulation (brush) of the hindpaw plantar surface, and then confirmed with brief pulses (∼1s, 21mW/mm^2^) from a 470nm LED. A fiber optic cable connected to a 470nm LED source (ThorLabs) was used to provide optogenetic stimulation of the hindpaw; stimuli were delivered with 10ms pulse widths at ∼21mW/mm^2^ with 10s ISI for 100 trials.

Electrophysiology data was amplified and bandpass filtered at 0.1-3000 Hz using a WPI ISO-DAM8 and acquired by a CED Micro1401 data acquisition system. Data was sampled between 12-34 kHz. Optogenetic stimulation was tracked on the data acquisition system using TTL logic.

### Histology and Immunohistochemistry

Following acute electrophysiology and behavior, spinal cord samples were extracted from mice and placed in 4% formaldehyde in 1x PBS overnight in 4 °C. Tissue samples were then placed in 30% sucrose in 1x PBS for 3 days, until samples sank. Samples were then frozen in OCT using a mixture of dry ice and 70% ethanol. A cryostat was used to take 25um sections of spinal cord samples, and either stored in −80 °C or immediately used for staining. Slides were washed in 1x PBS, three times for 10 minutes and then incubated with primary antibody (1:250 Rabbit anti-CGRP, Millipore-Sigma, PC205L) in blocking buffer solution (1x PBS, 1% Bovine Serum Albumin, 0.1% Triton X-100) in 4C for 24 hours. After primary antibody incubation, samples were washed with 1x PBS three times for 10 minutes and then incubated for one hour with secondary antibody (1:500 Goat anti-Rabbit Alexa Fluor 635, ThermoFisher, A-31577) in blocking buffer solution at room temperature. Samples were then washed three more times in 1x PBS for 10 minutes prior to mounting with Fluoromount-G (Thermo Fisher Scientific). If samples were additionally stained with the fluorescent Nissl marker, Neurotrace (NT) Blue, they were incubated with 1:200 NT in 1x PBS for 10 minutes at room temperature prior to washing and mounting.

### Custom behavioral platform and stimuli

A custom 3D printed platform was developed for restraining mice and delivering stimuli. The platform consists of two pieces that are epoxied together; a base and a grated platform. The base component features three M6 counter-bore holes to fix the platform to an aluminum breadboard (Thorlabs, MB4560/M) and a separate rectangular hole to fit the grated floor. The grated floor consists of a 5×5 grid of 4mmx4mm holes with 1mm spacing (**Fig. S1**) that is epoxied into the main base component. Stimulation was delivered through the central hole in the grated floor via a ferrule attached to a fiber optic cable (Thorlabs, M98L01). Optogenetic stimulation was delivered using a 470nm LED source (Thorlabs, M470F3) controlled by a LED driver (Thorlabs, LEDD1B) that received a modulatory analog input. Tactile stimuli were delivered by attaching the fiber optic cable to a piezo-electric plate bender (Noliac, CMBP09) that was actuated with a piezo haptic driver (Texas Instruments, DRV2667). All aspects of the behavior were controlled by a Beaglebone black development board.

### Pain detection task

All mice were habituated for a minimum of 3 days to the behavioral platform using techniques adopted from a similar restraint methodology^14^. On the first day, mice were head-posted for 15 minutes. On the second day, mice were head-posted and hind paw restricted using flexible medical tape for 30 minutes. On the third day, mice were head-posted and hind paw restricted for 1 hour. Following habituation to water restriction (5 days maximum), mice were then introduced to the water spout. Once mice displayed a persistent desire to lick over the course of 1 hour, training on the detection behavior began. All sessions (training and testing) consisted of 360 randomly distributed trials; 20 trials per each of the 17 target optogenetic stimuli (0.14-5.45 mW/mm2) and a single tactile catch stimulus (one cycle of ∼120Hz sine wave). Each trial also had a randomly distributed ISIs (between 3-6s). Response windows were randomized between 0.8-1.5s for each training session, while testing sessions maintained a 1s response window. Additionally, each subsequent day of training, the chance for receiving a water reward (5-10uL) on a hit trial was reduced by 5-10% until the reward chance reached 80%, which was the reward chance for all behavioral testing sessions. False alarm warning optogenetic pulses were set to 5.45 mW/mm2 at 10ms pulse widths with 100ms ISIs. Timeout’s were given whenever mice licked prior to the start of the delivery of a stimulus and lasted the length of the initial ISI for that trial.

### Video capture

Video data was acquired at either 60 or 120 fps, 1080p using an offline GoPro Hero6 Black and stored as mp4 files directly onto a microSD card. Triggers were recorded as analog input through the on-board audio channel and later extracted from the corresponding mp4 file in MATLAB.

### Acute electrophysiology data analysis

Data was analyzed in MATLAB using available packages and custom scripts. Electrophysiology data was bandpass (0.1-100Hz) and notch filtered with EEGLAB 9.0^20^ before data was down sampled to 8.3kHz. Data was then epoched with respect to the stimulus onset and averaged for each individual subject. Responses shown in **Fig. 1b** are normalized to the absolute maximal response in the averaged epoch.

### Behavioral data analysis

The sensitivity (d’) was used as a metric to determine which behavioral sessions were used for analysis. Calculation of d’ was performed as previously described^21^ for detection behaviors in mice. For each session considered, d’ was calculated in a 50-trial sliding window, every trial using the equation:

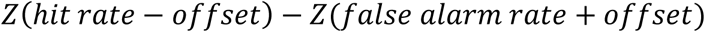

Where Z indicates the inverse of the Gaussian distribution, and the offset is set to 0.001 to avoid saturation of d’. Criterion was likewise calculated, using the equation: 

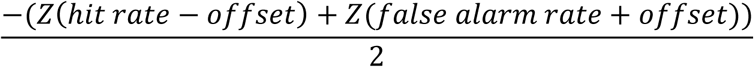

Psychometric responses from lick report and reflex report were fitted with a cumulative Gaussian function using the MATLAB toolbox psignifit 4^22^. Fitting was performed only on optogenetic stimulation trials, while tactile trials were shown as a fixed rate. Euclidean distance was calculated using the following equation:

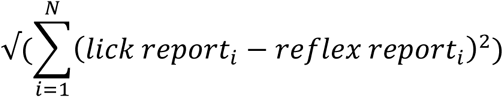

Euclidean distance was calculated for each stimulus intensity for each session (20 trials per intensity). Dashed similarity lines in **Fig. 2f** were calculated by determining the Euclidean distance to achieve 95% similarity (19 trials) and 50% similarity (10 trials) over a total of 20 trials.

### Video data analysis

Video data was distributed among 3 scorers who were blinded to both the animal, session number, and trial types. Scorers were on a 0-3 scale comparing pre-stimulus to post-stimulus movement, and criterion for scoring were decided prior to scoring. Scores of “0” indicated no change in movement. Scores of “1” indicated subtle movement, such as a shift in weight on the hindquarters, but no lifting of the hindquarters. Scores of “2” indicated a mild response where the mouse’s hindquarters raised a small amount, with slow movement. Scores of “3” indicated an overt response where movements were immediate and rapid, with either tail flicking, and raising of the hindquarters.

A custom MATLAB script was used to epoch each video trial to 250ms pre-stimulus to 500ms post-stimulus. Scorer’s were trained on a practice dataset (37 trials from an unused behavioral session, consisting of all trial types) prior to scoring behavioral data used for analysis.

### Statistical analyses

All statistical analyses were performed using Matlab R2019a. Data were tested for normality using a one-sample Kolmogorov-Smirnov test prior to any comparisons. In all comparisons between lick and reflex report, as well as comparisons made with the false alarm rates, significance was determined using a two-sided Wilcoxon signed rank test with *P* < 0.025. Multiple comparisons were performed using the Bonferroni correction. A one-sided Mann-Whitney U test was used to compare differences in overall hit rate across sessions between control and test sessions, with a significance level of *P* < 0.05.

## Data & code availability

Custom components developed for the behavior are available for download on github (https://github.com/neuromotion/nociceptive-detection). The data that support the findings of this study are available on request from the corresponding author.

## Acknowledgements

We would like to thank Stephanie Jones, Christopher Moore, Christopher Deister, and Hyeyoung Shin for advice and feedback on behavioral training, analysis, and experimental design. We would also like to thank Jacqueline Hynes for discussions on statistical analysis.

## Author Contributions

C.B., D.B. and C.S. conceived the experiments. C.B. designed and performed the experiments. C.B. performed all headpost and acute spinal electrophysiology surgeries, and all behavioral analyses. C.B. and M.E. carried out water restriction procedures and habituated mice. C.B., K.B., K.V. performed reflex scoring on all video data. C.B. and A.A. performed all histology and immunohistochemistry. C.B., C.S., and D.B. wrote the manuscript.

