## Supplemental Figures for "Automated and rapid self-report of nociception in transgenic mice"

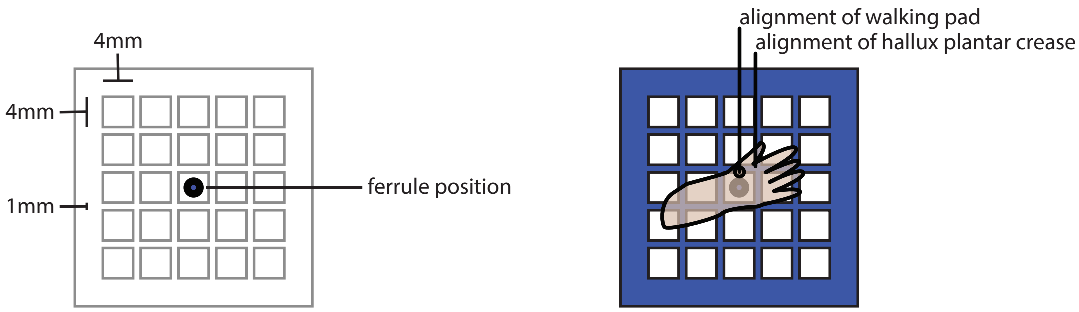


**Supplementary Figure 1.** Alignment of right hindpaw over 5x5 (4mm x 4mm holes with 1mm spacing) grated floor during behavior. Crease between the first and second digit on the left is aligned with respect to the corner square containing the ceramnic ferrule. Walking pads closest to the side of the foot were also used to position the hindpaw in place.

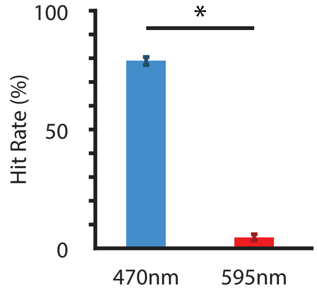


**Supplementary Figure 2.** Comparison of hit rate for 470nm LED (blue, n = 28 sessions) and 590nm control LED (red, n = 12 sessions), mean±SEM., significance of $P < 0.05$ using the Mann-Whitney U test.

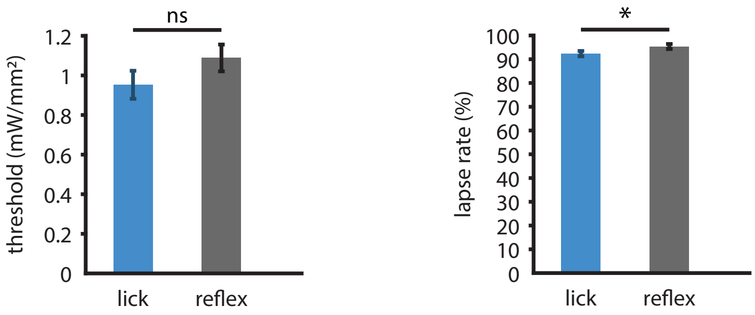


**Supplementary Figure 3.** Differences between extrapolated threshold and lapse rates between lick and reflex reports using psignifit matlab toolbox. N = 40 sessions from 5 mice, plotting mean±SEM, significance of $P < 0.025$ using the Wilcoxon signed rank test, multiple comparisons performed using the Bonferroni correction.

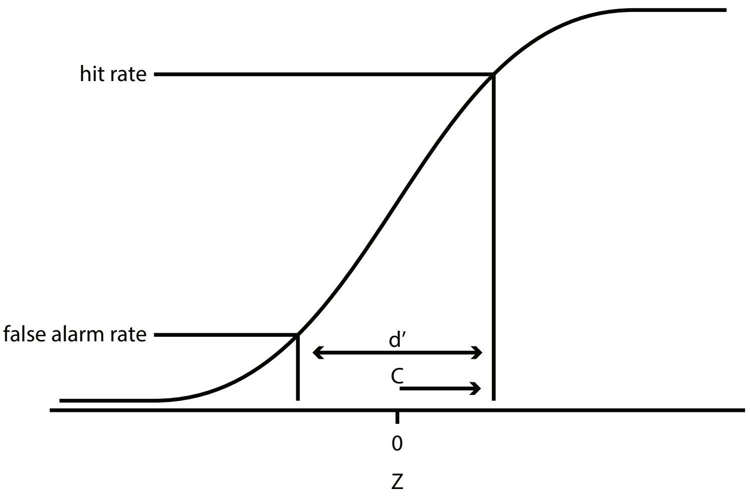


**Supplementary Figure 4.** Equations and illustration for calculating sensitivity (d’ ) and response bias (C) measures. While d’ is the difference between the signal and the signal + noise, it can be estimated by the difference of the inverse cumulative distribution function between the hit and the false alarm rates, whereas C can be estimated as one-half the sum of the inverse cumulative distribution function between the hit and false alarm rates. C will always be greater than 0, as 0 indicates the animals response biased to always reporting a stimulus.

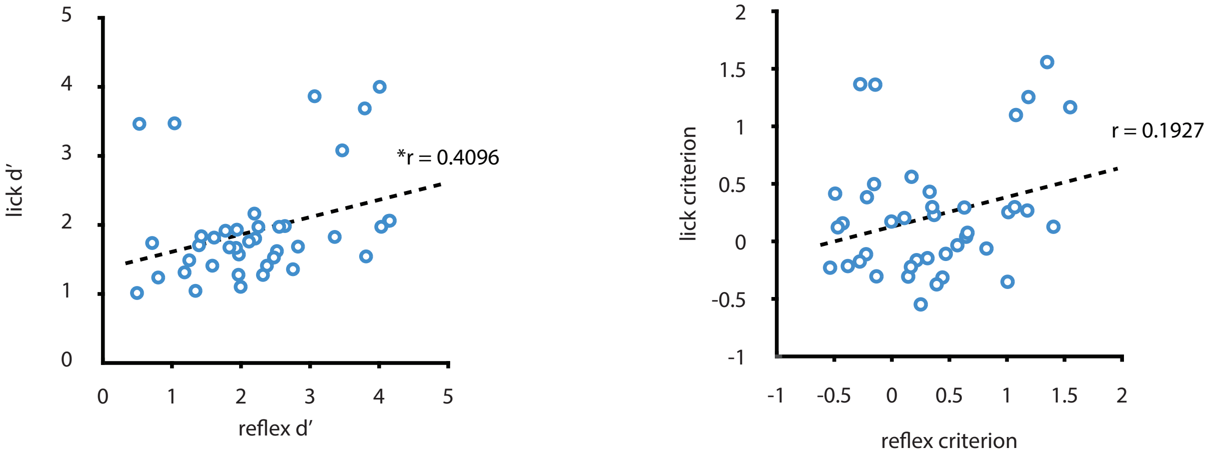


**Supplementary Figure 5.** Spearman's correlation of d' and criterion between lick and reflex for each session (n = 40).

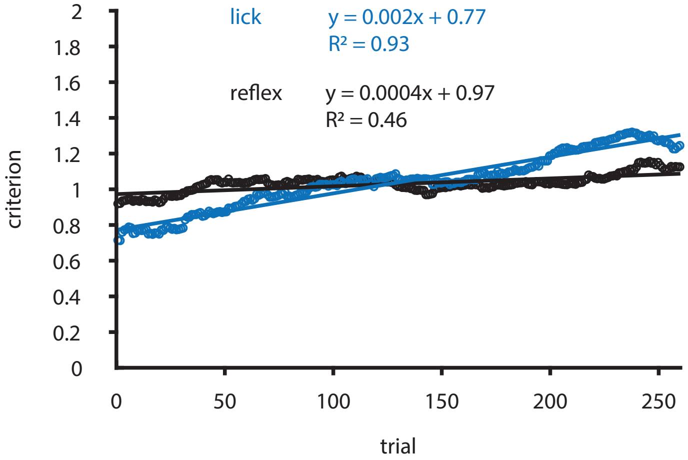


**Supplementary Figure 6.** Linear regression for lick and reflex criterion within sessions (n = 40). While both fits have a positive slope, the lick criterion has both a greater slope and $R^2$ than the reflex criterion, indicated a larger modulation of lick criterion over time.
